# Ooids forming *in situ* within microbial mats (Kiritimati atoll, central Pacific)

**DOI:** 10.1101/2021.05.05.442839

**Authors:** Pablo Suarez-Gonzalez, Joachim Reitner

## Abstract

Ooids (subspherical particles with a laminated cortex growing around a nucleus) are ubiquitous in the geological record since the Archean and have been widely studied for more than two centuries. However, various questions about them remain open, particularly about the role of microbial communities and organic matter in their formation and development. Although ooids typically occur rolling around in agitated waters, here we describe for the first time aragonite ooids forming statically within microbial mats from hypersaline ponds of Kiritimati (Kiribati, central Pacific). Subspherical particles had been previously observed in these mats and classified as spherulites, but they grow around autochthonous micritic nuclei, and many of them have laminated cortices, with alternating radial fibrous laminae and micritic laminae. Thus, they are compatible with the definition of ‘ooid’ and are in fact identical to many modern and fossil examples. Kiritimati ooids are more abundant and developed in some ponds and in some particular layers of the microbial mats, which has led to the discussion and interpretation of their formation processes as product of mat evolution, through a combination of organic and environmental factors. Radial fibrous laminae are formed during periods of increased supersaturation, either by metabolic or environmental processes. Micritic laminae are formed in closer association with the mat exopolymer (EPS) matrix, probably during periods of lower supersaturation and/or stronger EPS degradation. Therefore, this study represents a step forward in the understanding of ooid development as influenced by microbial communities, providing a useful analogue for explaining similar fossil ooids.

## Introduction

Ooids (subspherical particles with a laminated cortex growing around a nucleus, cf. Richter, 1983a) have fascinated and intrigued humanity for millennia (Burne et al., 2012; Weber, 2014). Despite being one of the sedimentary particles with the oldest continuous geological record, since the Archean (e.g. Siahi et al., 2017; Flannery et al., 2019) and with the longest history of descriptions and interpretations, since Roman times (Burne et al., 2012), they have prompted continuous discussions about their definition, classification, formation processes, mineralogy, diagenesis and evolution throughout Earth history (see some previous reviews in Kalkowsky, 1908; Bucher, 1918; Bathurst, 1968; Teichert, 1970; Fabricius, 1977; Davies et al., 1978; Simone, 1981; Krumbein, 1983; Richter et al., 1983a; Wilkinson et al., 1985). Some of these discussions are still active nowadays (see a recent review in Diaz and Eberli, 2019), mainly about determining the exact processes behind the origin and development of ooids. Being characteristic particles of active shoals in shallow agitated waters, they have been long considered physicochemical precipitates formed by constant rolling in the water (e.g. Duguid et al., 2010; Trower et al., 2018). Nevertheless, ooids have an equally long history of being interpreted as formed by some degree of influence from organic molecules or even microbial communities (e.g. Kalkowsky, 1908; Mitterer, 1968; Suess and Fütterer, 1972; Reitner et al., 1997; Diaz et al., 2017; Li et al., 2017; Mariotti et al., 2018). The work presented here entails a step forward in the knowledge of biotic factors on the origin of ooids, since it describes in detail for the first time ooids occurring within thick microbial mats of hypersaline ponds from Kiritimati atoll (Republic of Kiribati, Central Pacific). The aims of the study are to investigate if the analysed particles are compatible with the definition of ooids and if they grow directly within the microbial mats, a situation that has previously been only rarely and locally described (Friedmann et al., 1973; 1985; Krumbein and Cohen, 1974; Krumbein, 1983; Gerdes et al., 1994; 2000; Hubert et al., 2018), and which is poorly understood. Consequently, this study will also aim to interpret the biotic and environmental factors that may control ooid development within microbial mats, providing a useful modern analogue for explaining the origin of fossil ooids, especially of those whose origin is suspected to be related with benthic microbial communities (e.g. Kalkowsky, 1908; Krumbein, 1983; Neuweiler, 1993; Li et al., 2017; Antoshkina et al., 2020; Zwicker et al., 2020)

## General setting and materials

Located in the central Pacific and close to the Equator (1°55’ N, 157°25’ W), the island of Kiritimati (formerly Christmas Island) is the largest atoll on Earth, with a land surface area of ~360 km^2^, and the largest island of the Republic of Kiribati (Fig. 1; Valencia, 1977; Schoonmaker et al., 1985). The surface of the island shows a reticulate pattern made up of ~500 small and very shallow ponds (most of them <1 km wide and <2 m deep, Helfrich et al., 1973; Valencia, 1977) with salinities ranging from brackish to hypersaline (Fig. 1; Schoonmaker et al., 1985; Saenger et al., 2006). In most of them, cm- to dm-thick microbial mats develop covering the pond bottom (Fig. 2; Trichet et al., 2001; Arp et al., 2012; Schneider et al. 2013; Ionescu et al, 2015). The ponds are surrounded by sparsely vegetated areas of carbonate debris from the atoll substrate, mainly mollusc and coral fragments (Saenger et al., 2006; Arp et al., 2012). Kiritimati has an arid climate controlled by the El Niño-Southern Oscillation (ENSO), which causes significant variations in rainfall, ranging from dry periods with <200 mm annually, to humid periods with up to 3000 mm annually (Helfrich et al., 1973; Saenger et al., 2006; Morrison and Woodroffe, 2009; Arp et al., 2012). This contrast between dry and humid periods causes strong variations in the water level of ponds (up to 2.5 m, Helfrich et al., 1973) and in their salinities, which can be up to 6 times higher, when comparing general salinity ranges given by Helfrich et al. (1973) and Schoonmaker et al., (1985). In addition, most ponds are hydrologically closed systems and differences in water level of up to 1.2 m have been observed even in immediately adjacent ponds (Helfrich et al., 1973).

**Figure 1:**
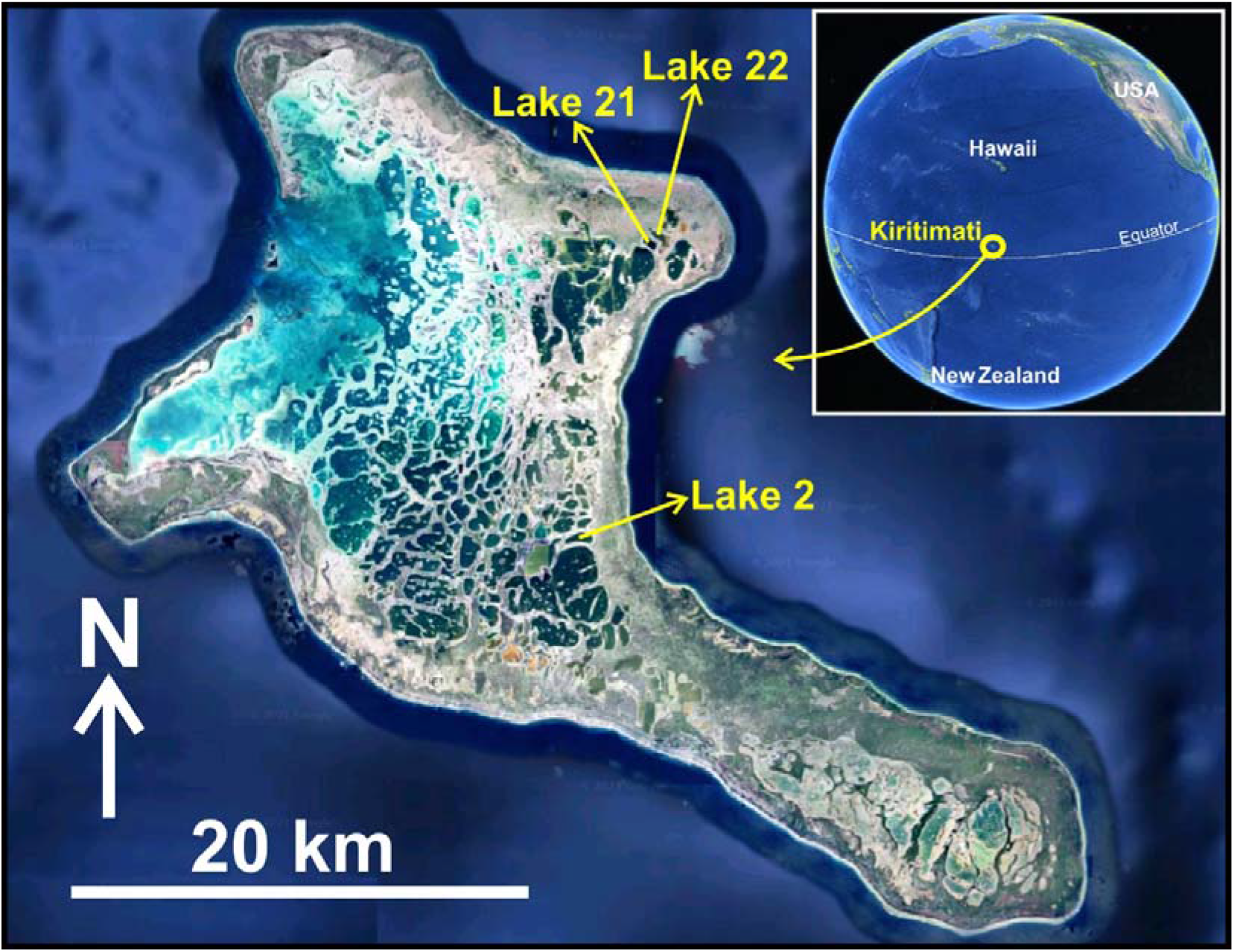
Satellite image from Google Earth of Kiritimati atoll, showing its reticulate pattern of ~500 small and shallow ponds, and highlighting those whose microbial mats have been studied here: Lakes 2, 21 and 22. Inset marks the location of Kiritimati in the central Pacific.

**Figure 2:**
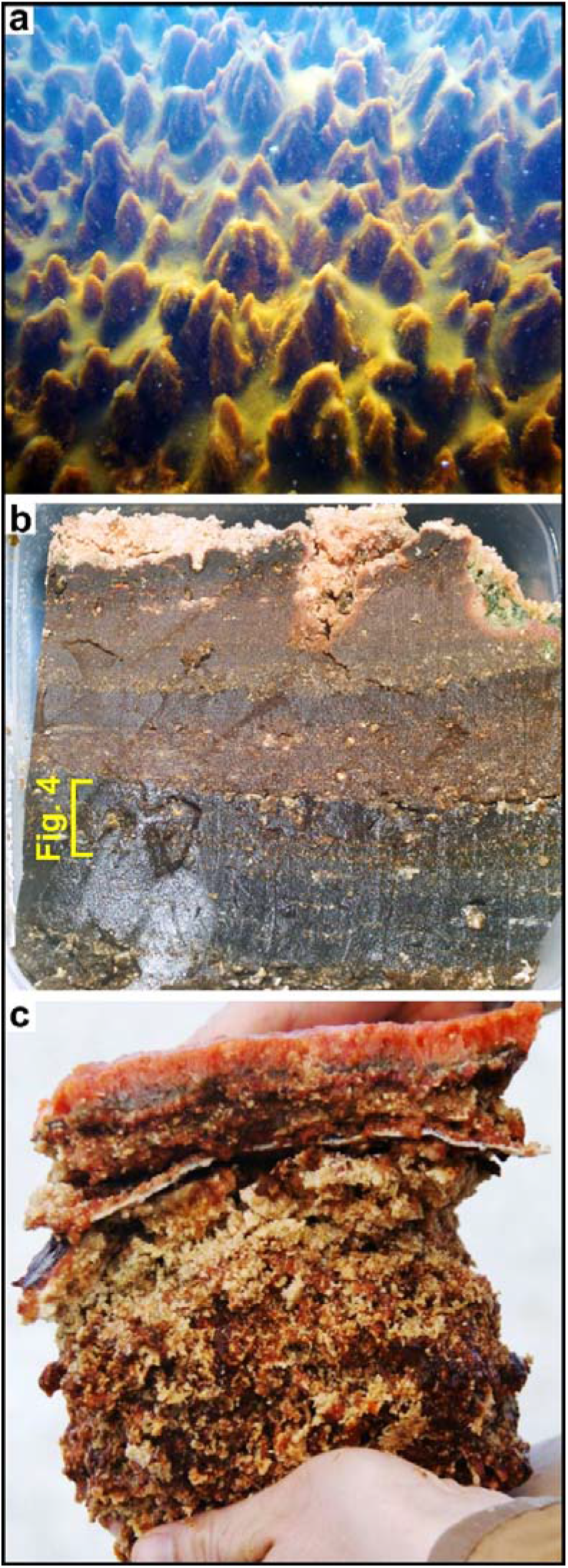
Samples studied in this work. **a** Subaqueous picture of the microbial mat covering the bottom of Lake 21, with conical protuberances, up to 10 cm tall. One of the protuberances was sampled and is studied here. **b** Freshly cut section of the microbial mat of Lake 22, ~12 cm thick, with its layers shown by colour banding. Lighter spots are mineral precipitates. The bracket marks the approximate location of the subsample whose photomicrograph is shown in Fig. 4. **c** Freshly cut section of the microbial mat of Lake 2. Note the fresh mucous exopolymers (EPS) of the photosynthetically active top layers contrasting with the lower more degraded layers. Lighter spots are mineral precipitates.

For this research, microbial mats from three different ponds (Lakes 2, 21 and 22; Fig. 1) were studied, all of them including abundant and active mineral precipitation. The sample from Lake 21 was taken from a central area of the pond, at ~1.5 m depth, and corresponds to the photosynthetically active uppermost ~8 cm of an orange-to green-coloured mat with conical protuberances and pinnacles (Fig. 2a; cf. Arp et al., 2012; Ionescu et al., 2015), whereas samples from Lakes 2 and 22 correspond to older and brownish microbial mats (~10 cm and ~12 cm thick, respectively), with flat tops, faint internal colour-layering, and with only the uppermost layer being photosynthetically active (Fig. 2b, c; cf. Blumenberg et al., 2015; Shen et al. 2018; 2020). The sample of Lake 2 was taken from a central area of the pond, at ~4 m depth (Blumenberg et al., 2015; Shen et al., 2020), whereas that of Lake 22 was taken from the pond shore, close to the mouth of a small, dry, ephemeral creek flowing into the pond (Shen et al., 2018). Previous ^14^C dating of the Lakes 2 and 22 mats have provided ages from 62 ± 40 years BP at the top and 1,291 ± 40 years BP at the bottom (in Lake 22, Shen et al., 2018) and from 62 ± 40 years BP at the top and 1,440 ± 40 years BP at the bottom (in Lake 2, Blumenberg et al., 2015).

All studied microbial mats consist predominantly of a gelatinous organic matrix (mainly formed by the exopolymers-EPS-secreted by the microbes) with mineral particles within it (Fig. 2b, c). Consistency and thickness of the organic matrix, as well as the amount of mineral precipitates varies through each mat and between different mats. Typically, upper layers of the mats show an abundant, fresh, and firm gelatinous matrix, with few minerals, whereas lower layers of the mats are crumblier due to less abundant and more degraded organic matrix and to larger and more abundant mineral precipitates (Fig. 2c). Nevertheless, local variations in mineral abundance, not following the downwards increase, are also observed between adjacent mat layers (Fig. 2b). The mineralogy of the precipitates is mainly aragonite, with gypsum occurring in some layers of the mats, typically at the top, and with minor traces of Mg-calcite, halite and protodolomite (Arp et al., 2012; Suarez-Gonzalez et al., 2017; Ionescu et al 2015; Shen et al., 2020). Two main types of aragonitic mineral precipitates occur within the mats: micritic aggregates and subspherical particles, which are the focus of this study and will be described in detail below. The micritic aggregates have a micropeloidal texture and range from mm-scale irregular aggregates in the upper parts of the mats, to cm-scale lumps with reticular structure downwards in the mats (Défarge et al., 1996; Trichet et al., 2001; Arp et al., 2012, Suarez-Gonzalez et al., 2017). Similarly, subspherical particles are typically smaller and less abundant in the upper parts of the mats, and larger and more abundant downwards, commonly coalescing with each other through micritic patches and bridges (Arp et al., 2012; Schneider et al., 2013; Suarez-Gonzalez et al., 2017).

## Methods

Sampling was conducted in 2011 and all samples were kept at −20°C until laboratory preparation. From each mat, several correlative adjacent histological thin sections were prepared covering the whole mat thickness. Samples were dehydrated with graded ethanol and embedded in LR White resin (London Resin Company Ltd., Reading, UK). Embedded samples were cut to a ~100 μm thickness using a microtome saw (Leica SP1600) and mounted on glass slides with Biomount mountant (Electron Microscopy Sciences, Hatfield, PA). Thin sections were observed under petrographic (Zeiss Axiolab) and fluorescence (Zeiss Axio Imager Z1) microscopes. In addition, mineral particles were separated from their organic matrix for their study in the electron microscope. Organic matter of the samples was oxidized with 6% NaOCl, changing the solution every 12 h (Mikutta et al., 2005) until traces of organic matter were no longer visible. Mineral particles were washed with distilled H_2_O until neutral pH was reached, and then dried. Some of the subspherical particles focus of this study were mechanically broken to observe their internal structure. The particles were sputtered with Pt/Pd (14.1 nm for 5 min) and observed in a field-emission scanning electron microscope (FE-SEM) Leica EM QSG100, using a detector of secondary electrons (SE2) at a voltage from 2 to 4 kV, combined with an INCA X-act energy dispersive X-ray (EDX) spectroscope (Oxford Instruments). Some histological thin sections were also studied with SEM and they were previously etched by submerging them for 10-30 seconds in a 5% EDTA (ethylenediaminetetraacetic acid) solution, for a better observation of the internal structure of mineral particles.

## Note on terminology

The scientific literature about subspherical carbonate particles and coated grains dates back for more than a century, with ongoing discussions and contrasting definitions (e.g. Peryt, 1983; Richter, 1983a; 1983b). Therefore, it is advisable to specify beforehand the classifications and definitions that will be used in this study. The main terms that will be applied to the subspherical particles studied are ‘spherulites’ and ‘ooids’. A general crystallographic approach to ‘spherulites’ defines them as “radially polycrystalline aggregates with an outer spherical envelope” (Shtukenberg et al., 2012), whereas geological points of view emphasize their “radial internal structure arranged around one or more centers” and the fact that they are “formed in a sedimentary rock in the place where [they are] now found” (Bates and Jackson, 1980, in Verrechia et al., 1995). Concerning ‘ooids’, also a purely descriptive definition is adopted, following Richter (1983a), who emphasizes that they are subspherical particles “formed by a cortex and a nucleus variable in composition and size”, where “the cortex is smoothly laminated” with laminae typically concentric. Therefore, the main difference between both types of subspherical particles is that unlike spherulites, ooids grow around a nucleus and show internal lamination. Although ‘spherulites’ and ‘ooids’ may also be envisaged as end-members of a gradational continuum and, in fact, intermediate steps between them do occur (e.g. Friedmann et al., 1973; Kahle, 1974), their two clearly different descriptive definitions are adopted here, for avoiding confusions between them, as well as genetic implications.

## Description of subspherical particles

Subspherical particles have been observed in the three studied microbial mats, although with different features and abundances between them and between each mat layer. In general, they range from spherical to ellipsoidal in shape, and from 0.1 to 3 mm wide (Figs. 3-8). Their outer surface is smooth, as seen with a hand lens (Fig. 3a, b), but SEM imaging reveals that it is irregular in detail, often pitted (Fig. 3c-e). In addition, botryoidal or domal overgrowths that cover only partially the particle surface are also observed (Fig. 3d-e, 5). Subspherical particles occur throughout the microbial mats, but they are typically larger and more abundant downwards, although significant differences in their abundance are observed between adjacent mat layers (Figs. 2b, c, 4). In the young and fresh mat of Lake 21 only very small subspherical particles occur (Figs. 3c, d, 5c), whereas larger ones are observed in the thicker and older mats of Lakes 2 and 22 (Figs. 3a, b, d, 4, 6), especially in their lower parts, where some subspherical particles are merged together forming irregular aggregates up to several centimeters long (Fig. 3b).

**Figure 3:**
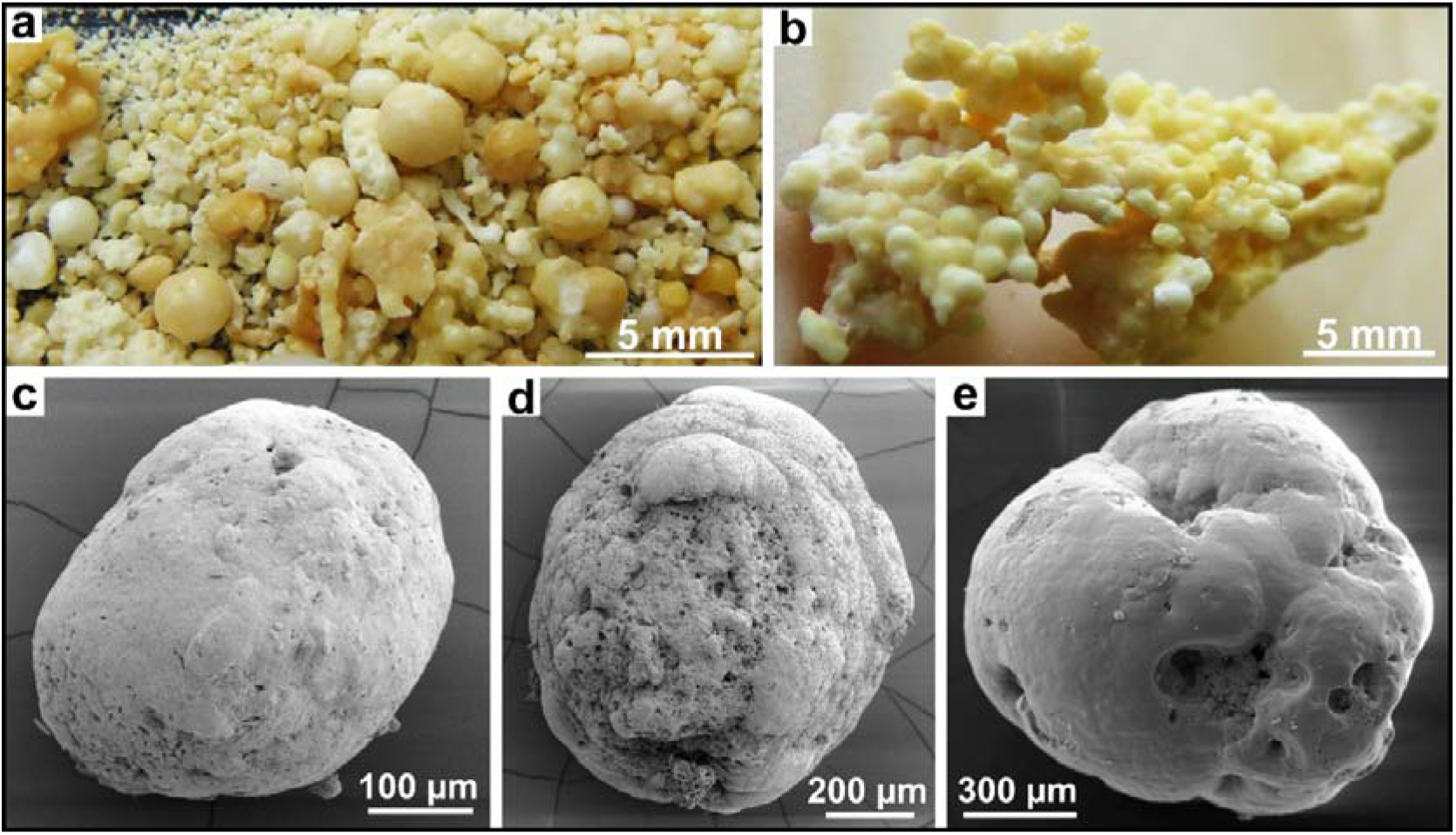
**a** Loose mineral precipitates from the upper part of the microbial mat of Lake 2 after removal of the organic matter. Note the abundance and diversity of sizes of subspherical particles. **b** Centimetric irregular aggregate from the lower part of the Lake 2 mat, formed by many subspherical particles merged together. **c-e** SEM images of subspherical particles, showing their irregular and often pitted surface, with botryoidal or domal overgrowths that cover only partially the particle. **c-d** from the Lake 21 mat, **e** from the Lake 2 mat.

**Figure 4:**
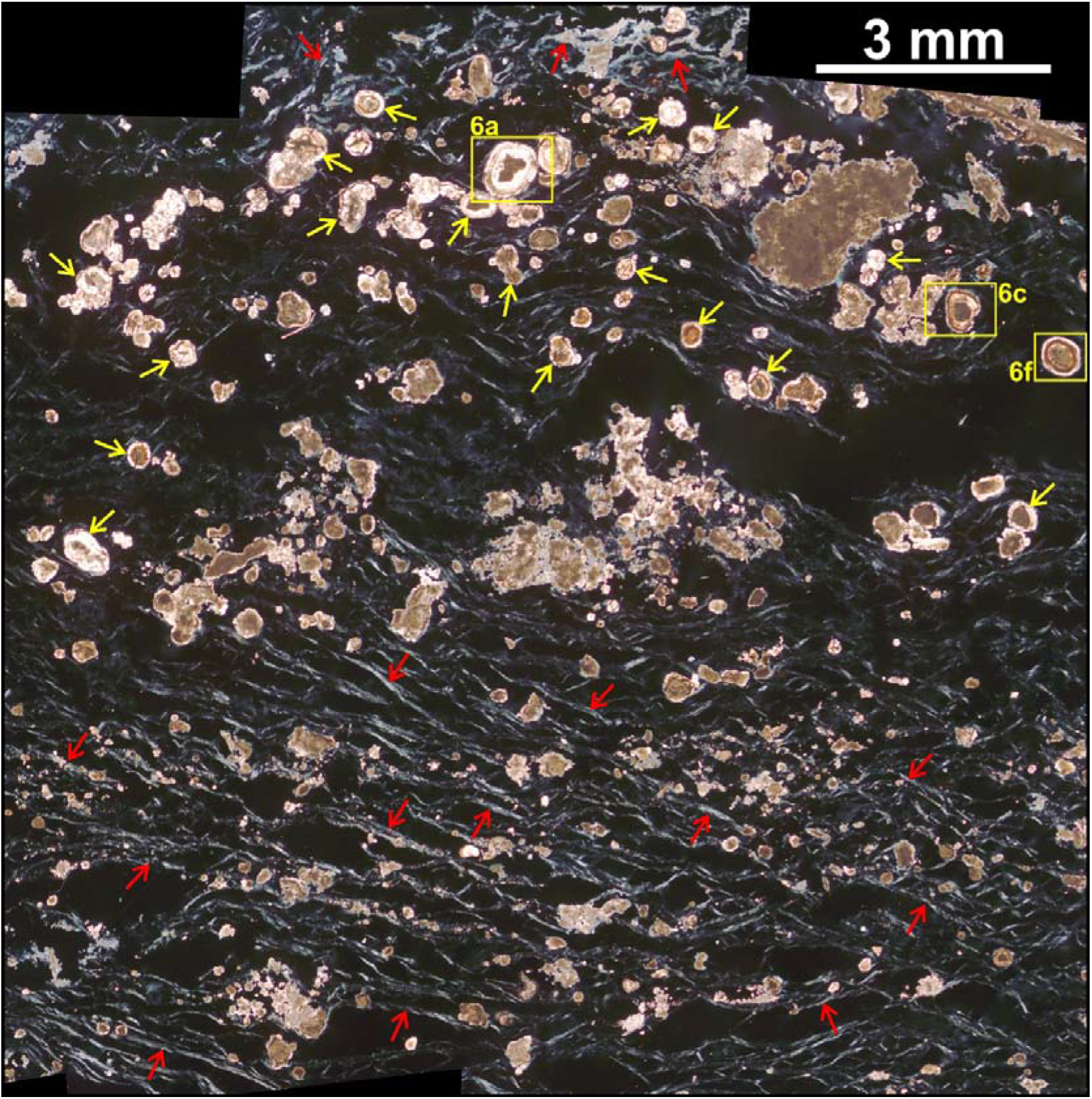
Crossed polarized light photomicrograph of a thin section from the lower middle part of the microbial mat of Lake 22 (see location in Fig. 2b). This sample includes the characteristic aragonitic precipitates of the Kiritimati mats: irregular micritic aggregates with micropeloidal texture, occurring throughout the sample, and subspherical particles (yellow arrows), occurring exclusively in a particular level. Yellow rectangles mark the location of Figs. 6a, c and f. Note that the EPS matrix has a stronger birefringence (red arrows) in the levels above and below the level rich in subspherical particles. This birefringence is caused by the degradation of EPS molecules (Arp et al., 1998; 1999; Reitner et al., 2005).

**Figure 5:**
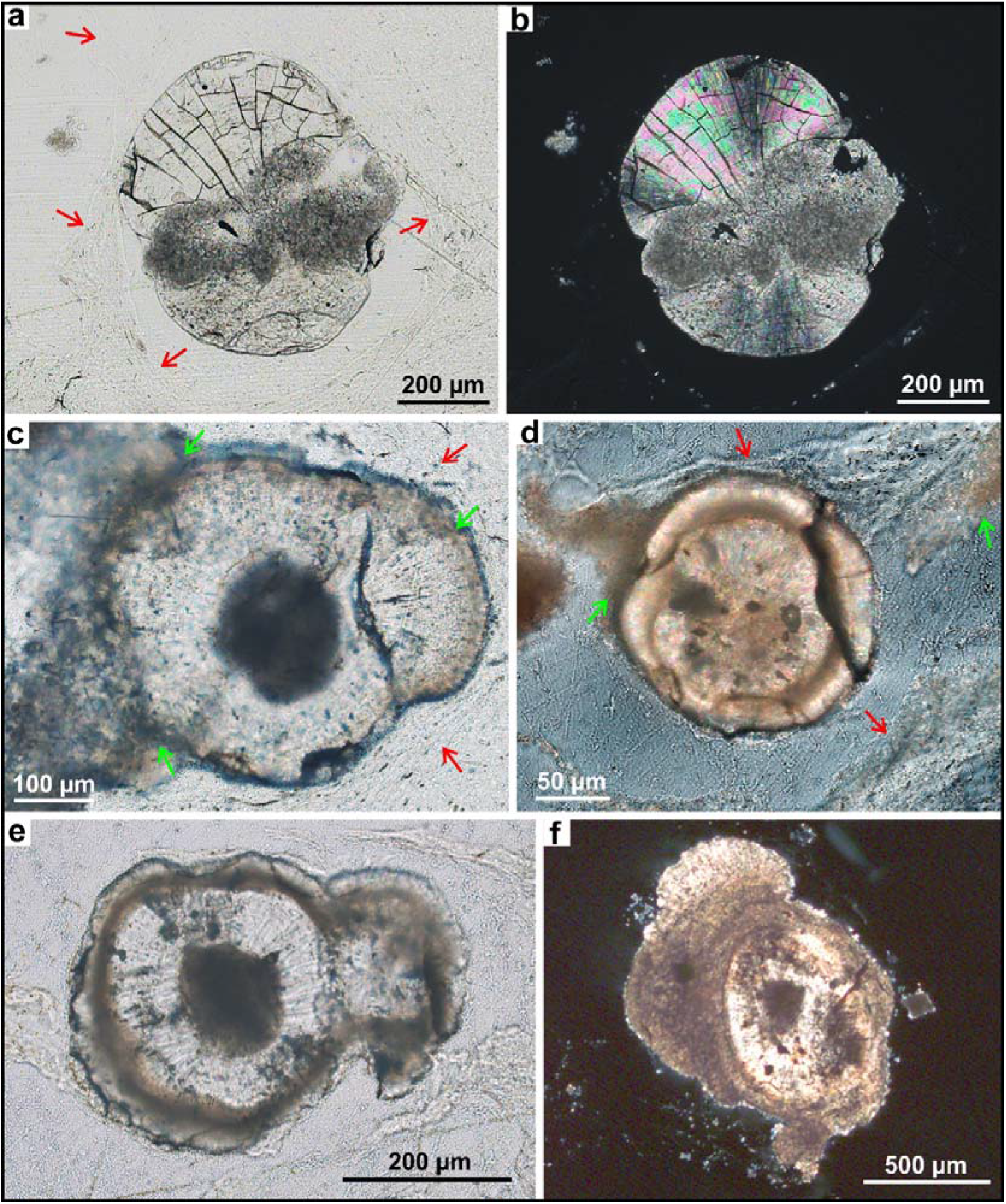
Early development stages of the Kiritimati subspherical particles. In some images the EPS matrix is slightly shrunk and detached from the particles due to the alcohol dehydration process during the preparation of thin sections (see Methods). **a** Transmitted light photomicrograph of a subspherical particle from the topmost layer of the Lake 2 mat, showing two tuft-like growths of radial fibrous aragonite above and below a nucleus of micropeloidal micrite. Red arrows point to threads of the EPS matrix surrounding the particle. **b** Same as **a** with crossed polarized light, which highlights the radial fibrous texture of the incipient cortex. **c** Transmitted light photomicrograph of a subspherical particle from the Lake 21 mat, showing incipient micritic laminae (green arrows) associated with the EPS matrix surrounding the particles (red arrows). **d** Crossed polarized photomicrograph of a subspherical particle from the Lake 22 mat, showing laminated cortex and micrite precipitation (green arrows) associated with the EPS matrix surrounding the particles (red arrows). **e** Transmitted light photomicrograph of two merged subspherical particles from the Lake 22 mat, showing an incipient laminated cortex, with a second radial fibrous lamina developing over the micritic lamina. **f** Crossed polarized photomicrograph of a complex subspherical particle from the Lake 2 mat, showing tuft-like micritic and radial fibrous overgrowths that do not cover completely the particle surface.

**Figure 6:**
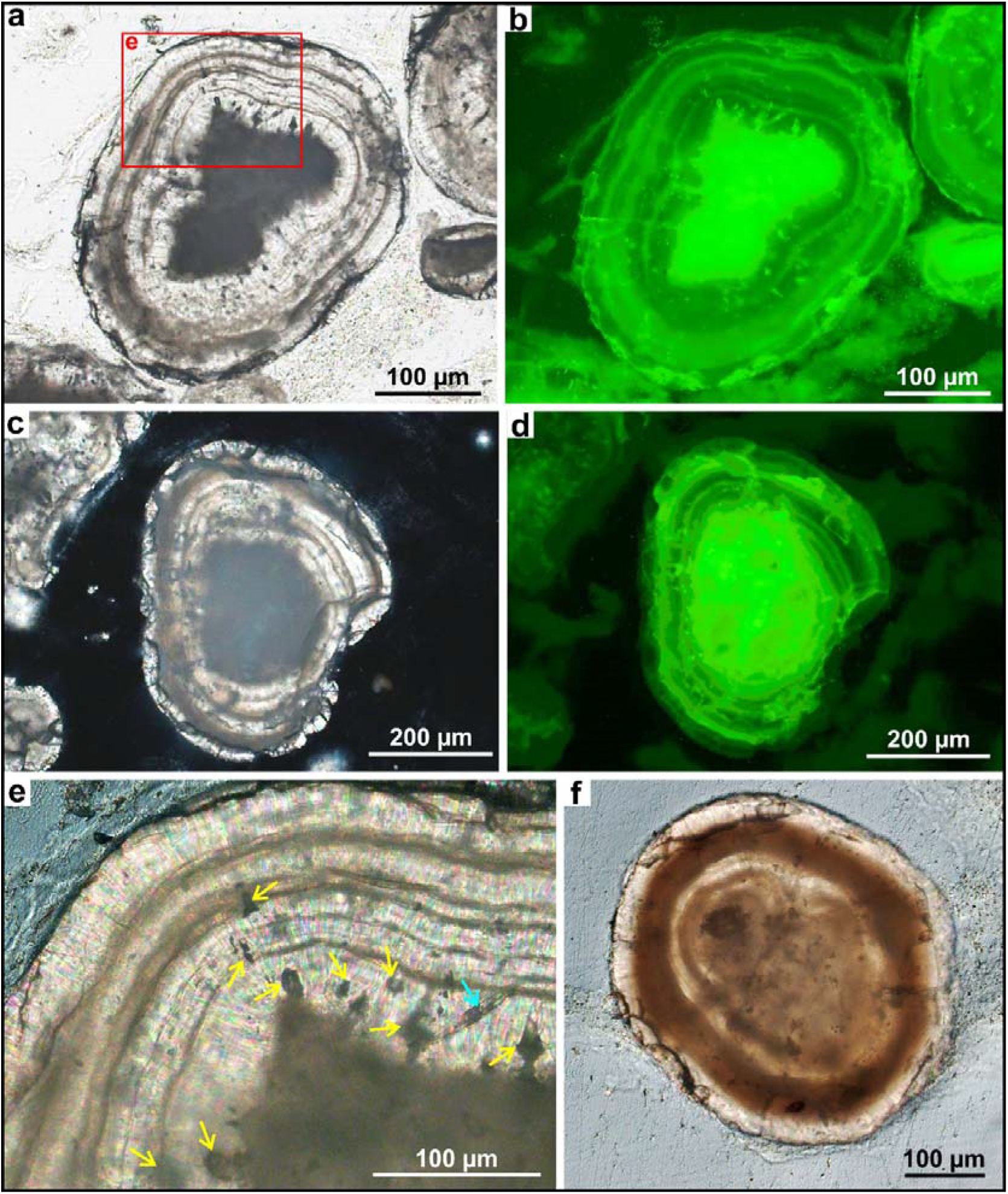
Subspherical particles with well developed laminated cortices (i.e. ooids) from the Lake 22 mat (see location in Fig. 4). **a, b** Coupled transmitted light and fluorescence photomicrographs of the same ooid. Red rectangle marks location of **e**. **c, d** Coupled crossed polarized and fluorescence photomicrographs of the same ooid. Note in **b** and **d** the stronger fluorescence of the nuclei and micritic laminae. **e** Crossed polarized photomicrograph from **a**. Yellow arrows point to dark inclusions within the cortex. Compare with **b** to note the stronger fluorescence of these inclusions. Blue arrow points to a diatom mold. **f** Transmitted light photomicrograph of and ooid with a thicker micritic lamina.

The internal structure of the subspherical particles consists of a nucleus and a cortex. The nuclei are always irregular micritic aggregates with micropeloidal texture, identical to the micritic aggregates that precipitate throughout the Kiritimati microbial mats (Figs. 4-6). SEM imaging shows that the nuclei consist of nm-scale aragonite crystals oriented randomly or forming μm-scale spherules, and with abundant EPS fibers and sheets between them and even some calcified microbe remains (Figs. 7, 8). The cortices of the particles have a radial fibrous structure formed by long and thin aragonite crystals, and some of the cortices have internal lamination and others do not (Figs. 5-8). The subspherical particles with laminated cortices are thus classifiable as ooids (*sensu* Richter, 1983a). Those with non-laminated cortices are thus closer to the definition of spherulites (*sensu* Bates and Jackson, 1980, in Verrechia et al., 1995), with the particularity that they do not grow around a “center” but around a micritic nucleus. Although subspherical particles occur throughout all the studied mats, those with well-developed laminated cortices seem to be especially abundant in particular layers (Fig. 4), typically at the lower and older parts of mats of Lakes 2 and 22, being absent from the mat of Lake 21.

**Figure 7:**
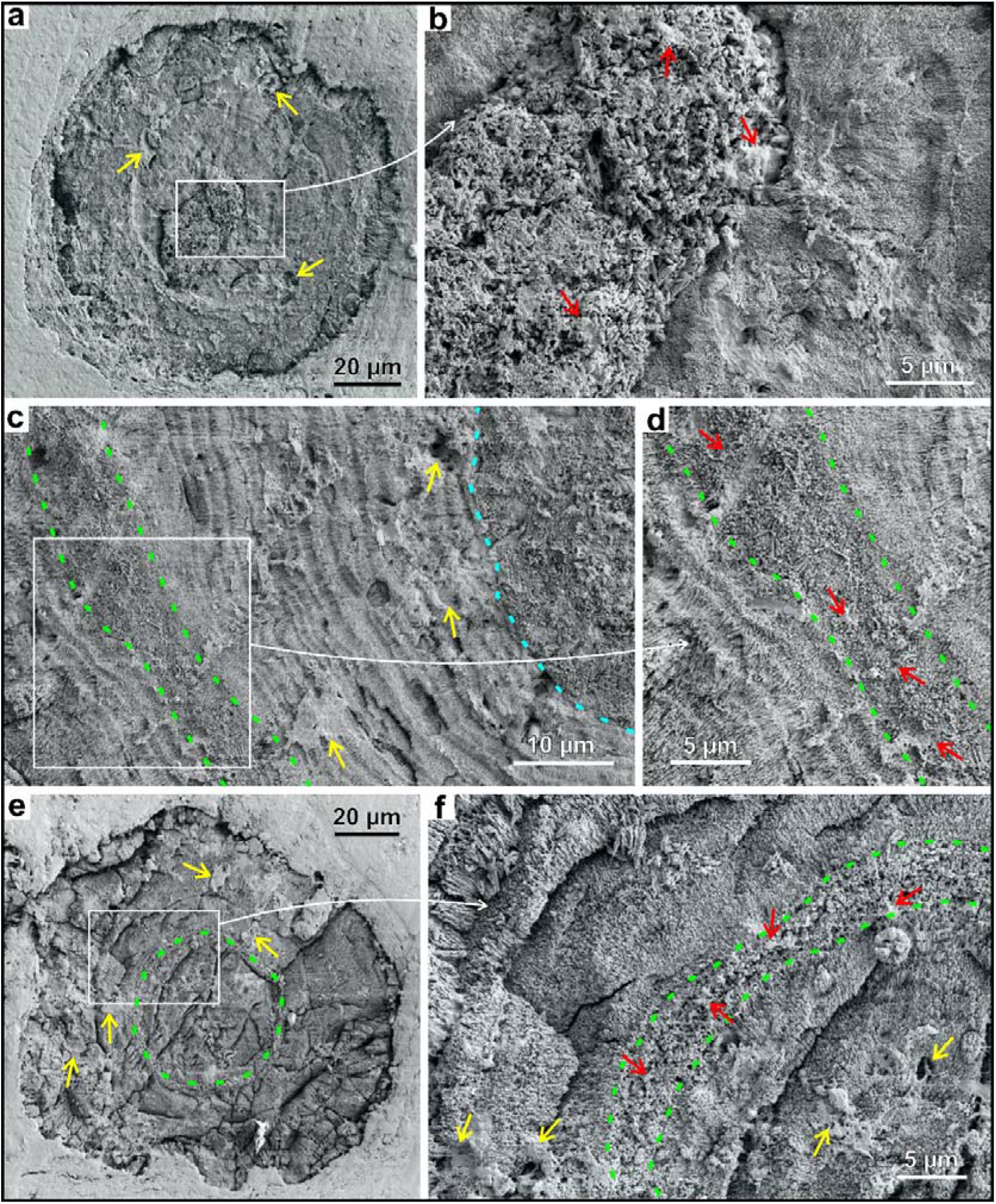
SEM pictures of EDTA-etched thin sections (see Methods) from the Lake 22 mat. **a** Ooid with a strong contrast between the micritic nucleus and the cortex. Yellow arrows point to cavities in the cortex filled with EPS. **b** Detail of **a**, showing the contact between the nucleus, formed by randomly oriented aragonite crystals surrounded by abundant EPS (red arrows), and the cortex, formed by radially oriented and elongated aragonite needles. **c** Detail of an ooid with micritic nucleus (blue dashed line marks its outer boundary) and a laminated cortex with radial fibrous laminae and a micritic lamina (highlighted with green dashed line). Note the finer lamination within the radial fibrous laminae. Yellow arrows point to cavities in the cortex filled with EPS. **d** Detail of **c**, showing the contrast between the radial fibrous laminae and the micritic lamina (bounded by the green dashed lines), which is composed of small and randomly oriented aragonite crystals surrounded by abundant EPS (red arrows). **e** Ooid with a micritic lamina highlighted by the green dashed line. Yellow arrows point to cavities in the cortex filled with EPS. **f** Detail of **e**, showing the contrast between the radial fibrous laminae and the micritic lamina (bounded by the green dashed lines), which is composed of small and randomly oriented aragonite crystals surrounded by abundant EPS (red arrows). Yellow arrows point to cavities in the cortex filled with EPS.

**Figure 8:**
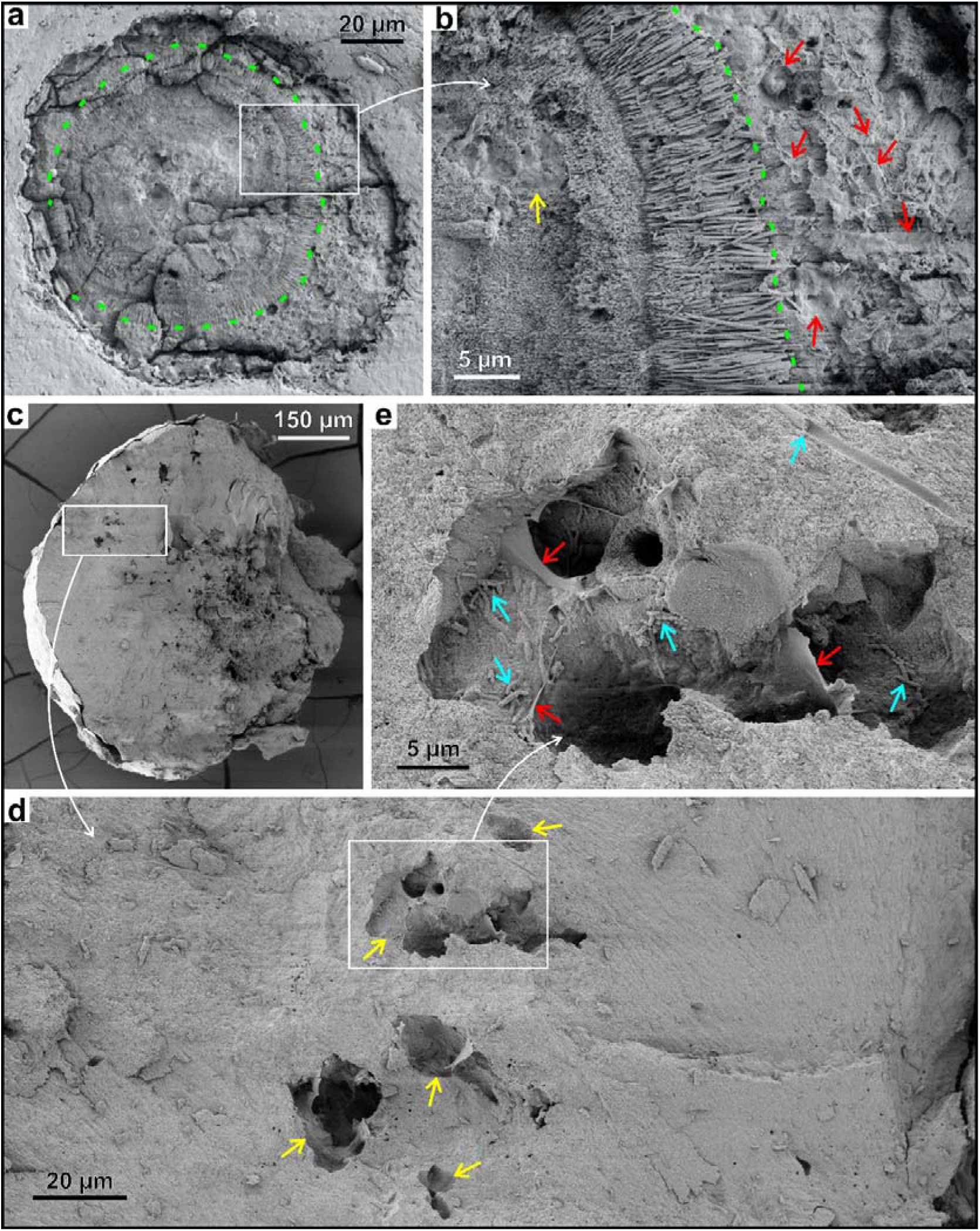
**a** SEM picture of an EDTA-etched thin section (see Methods) from the mat of Lake 22, showing an ooid with a micritic lamina forming at its external surface. Green dashed line marks the contact between the micritic lamina and the underlying radial fibrous lamina. **b** Detail of **a** showing the contact (highlighted with green dashed line) between the inner radial fibrous laminae and the outer micritic lamina, which is composed of small and randomly oriented aragonite crystals surrounded by abundant EPS (red arrows). Yellow arrows point to cavities in the cortex filled with EPS. **c** SEM picture showing the inside of a broken subspherical particle (from the mat of Lake 21), with its nucleus on the right side and the cortex on the left. **d** Detail of **c** showing several cavities in the cortex filled with EPS. **e** Detail of **d** showing the inside of one the cavities. Red arrows point to EPS inside the cavity. Blue arrows point to calcified microbe remains inside and outside the cavity.

In the particles with laminated cortices, their lamination is caused by thin micritic laminae that periodically interrupt the fibrous radial aragonite growth (Figs. 5-7). The thickness of micritic laminae is laterally variable, but they are typically only a few μm thick, locally reaching 60 μm (Figs. 5-8). Micritic laminae consistently show a stronger fluorescence than the adjacent fibrous radial laminae (Fig. 5b, d). Locally, micritic laminae occur not within the cortices, bounded by fibrous radial laminae, but at the external surface of subspherical particles, covering them partially or completely as their youngest lamina, and being associated with the EPS matrix that surrounds the particles (Fig. 5c, d, f, 8a, b). Contrasting with the long and radially oriented aragonite crystals of the fibrous radial laminae, the micritic laminae consist of nm-size aragonite crystals oriented randomly and with abundant EPS between them (Figs. 7c-f, 8a, b). In some particles, a finer lamination is also observed within the fibrous radial laminae, which are subdivided in 0.5-2 μm thick laminae that do not seem to interrupt the continuous growth of the aragonite crystals (Fig. 7c, d).

Regardless of whether they are internally laminated or not, most cortices of subspherical particles show dark inclusions up to 30 μm wide with strong fluorescence (Fig. 6a-e). These inclusions are cavities within the particle cortex, which include abundant EPS and some calcified microbe remains (Figs. 7, 8). In addition, calcified microbe remains, mainly filamentous bacteria and diatoms, are also locally observed enclosed by the fibrous aragonite crystals of the cortices and outside of the dark inclusions, especially in particles of the Lake 2 mat (Figs. 6e, 8e).

## Discussion

### *In situ* growth of ooids within microbial mats

Although subspherical particles of the Kiritimati microbial mats have been previously referred to as ‘spherulites’ (Défarge et al., 1996; Arp et al., 2012; Schneider et al., 2013; Ionescu et al., 2015) or ‘spherules’ (Schmitt et al., 2019; Chen et al., 2020), the detailed description presented here shows that some of the subspherical particles fit perfectly the definition of ‘ooids’ (Richter, 1983a), as they are composed of a laminated cortex growing around a nucleus. Those with non-laminated cortices might be classed as ‘spherulites with nucleus’ or ‘non-laminated ooids’ and are equivalent to other examples of modern ooids with non-laminated cortices, such as some ooids from Great Salt Lake (Eardley, 1938; Kahle, 1974; Reitner et al., 1997; Chidsey et al., 2015).

Independently of their terminological classification, the features of both the ooids *sensu stricto* and the non-laminated ooids, indicate that they were formed and developed directly within the studied microbial mats. Firstly, no equivalent particles have been observed, nor previously described, in or around the ponds of Kiritimati (Saenger et al., 2006; Arp et al., 2012) and, thus, they cannot be allochthonous particles transported to and trapped within the microbial mats of the ponds (cf. Suarez-Gonzalez et al., 2019). In addition, the subspherical particles grow around EPS-rich irregular micritic aggregates identical to the micritic aggregates that precipitate throughout the Kiritimati microbial mats (Figs. 4-8; Défarge et al., 1996; Trichet et al., 2001; Arp et al., 2012, Suarez-Gonzalez et al., 2017). Similarly, micritic laminae of the ooids show high fluorescence, due to their content in EPS (Fig. 6b, d), and both ooids and non-laminated ooids include abundant EPS-rich cavities and calcified microbe remains (Figs. 6e, 7, 8), all of them likely enclosed within the mineral structure during *in situ* growth of their cortices. The fact that ooids at different developmental stages occur within the mats, from ooids with incipient micritic laminae forming around them (Figs. 5, 8a, b) to fully-developed laminated ooids (Fig. 6, 7), which even coalesce with each other in their growth (Figs. 3b, 4, 5e), further supports their *in situ* origin. Other examples of ooids, very similar to those of Kiritimati, have been also described in microbial mats from shallow hypersaline settings and interpreted as formed *in situ* within the mats (Friedmann et al., 1973; 1985; Krumbein and Cohen, 1974; Krumbein, 1983; Gerdes et al., 1994; 2000; Hubert et al., 2018).

### Processes of ooid formation and development within microbial mats

The *in situ* growth of the Kiritimati ooids prompts evaluating the conditions that underlie their formation and the processes that allow their continuing development. Previous works studying the so-called ‘spherulites’ of Kiritimati (Défarge et al., 1996; Arp et al., 2012), have highlighted their recent occurrence already at the youngest photosynthetically active layer of the mats, together with their intimate association with the fresh EPS matrix of these layers, and their very positive δ^13^C values. These features have led to interpret them as early precipitates of the mats, formed through the combination of high aragonite supersaturation, caused by intense photosynthesis, and efficient inhibition of precipitation by pristine EPS (Arp et al., 2012). This combination of factors produces that radial fibrous aragonite precipitates only at few spots where inhibition is overcome and nucleation points exist, i.e. around preexisting micritic carbonate nuclei. This plausible explanation accounts, however, only for the first radial fibrous lamina of the particle cortex, not for the successive alternation of radial fibrous and micritic laminae that forms the characteristic lamination of the ooids described here. In other examples of ooids growing within hypersaline microbial mats, their lamination has been explained as caused by biologically induced chemical changes within microenvironments of the mat, which produce alternation of EPS-rich dark laminae and lighter fibrous aragonite laminae (Gerdes et al., 1994).

In the case of the Kiritimati ooids, the fact that their abundance and development vary not only in different layers of the same mat but also between mats of different ponds (Figs. 2-4), suggests that besides the biological influence, probably also environmental factors are involved in ooid development. The young and photosynthetically active mat of Lake 21 includes only small non-laminated ooids with a single radial fibrous aragonite lamina (Fig. 5), in agreement with the interpretation of Arp et al. (2012) of photosynthesis-induced high supersaturation coupled with strong EPS inhibition of precipitation. The older and more layered mats of Lakes 2 and 22 contain laminated ooids, which are generally larger and more abundant downwards, indicating that the lamination caused by alternation of radial fibrous laminae and micritic laminae is developed during the burial of older mat layers under overlying younger ones. The fact that radial fibrous laminae include EPS-filled cavities and microbe molds enclosed by the aragonite crystals (Figs. 6e, 7, 8) suggests that the precipitation of these laminae may have been relatively rapid and episodic, entombing mat remains that are very well preserved. This is consistent with the interpretation of precipitation occurring only locally and occasionally when very high supersaturation overcomes strong inhibition by fresh EPS. On the other hand, micritic laminae observed forming at the outer surface of ooids are intimately associated with the EPS surrounding the ooid (Figs. 5, 8a, b), and micritic laminae within the ooid cortices include EPS remains between the aragonite crystals (Fig. 7d, f). These facts indicate that the precipitation of micritic laminae is more directly and strongly controlled by the EPS matrix of the mat, which likely caused a slower precipitation of more abundant but much smaller and irregularly oriented aragonite crystals (as compared to the radial fibrous laminae), probably triggered by lower supersaturation and/or weaker inhibition due to increasing degradation of EPS with burial (cf. Trichet and Défarge, 1995; Reitner et al., 1995; Baumgartner et al., 2006; Arp et al., 2012).

The repeated alternation of radial fibrous and micritic laminae that forms ooid cortices indicates that both precipitation mechanisms interpreted above (rapid fibrous radial precipitation and slower and more EPS-controlled micrite precipitation) alternated successively in the same space. Since back-and-forth variations on EPS inhibition of precipitation are unlikely because decreasing inhibition is due to the progressive degradation of EPS with burial, it is more plausible that the alternation of both precipitation mechanisms is driven by variations in aragonite supersaturation within the mat microenvironment. Such variations can be caused by biotically-influenced changes within the mat and/or by changes in the overall pond hydrochemistry. The case of the Lake 22 is particularly relevant, because it includes the most abundant ooids with well-developed laminated cortices, and because these are especially concentrated in one particular layer at the lower middle part of the mat (Figs. 2, 4). Interestingly, the ooid-rich layer also shows a marked difference in the EPS matrix, when compared with the matrices of its adjacent layers, which are very birefringent under crossed-polarized light (Fig. 4). This birefringence is caused by a significant sugar-rich EPS degradation (Arp et al., 1998; 1999; Reitner et al., 2005), and thus the EPS matrix of the ooid-rich layer seems to be less degraded than the layers above and below, which include abundant irregular micritic aggregates but hardly any ooids (Fig. 4). This points to a probable relationship between the development of ooids and a stronger inhibition effect due to less degraded EPS. In addition, the mat of Lake 22 differs from the others in that it was sampled at the shore of the pond and by a small dried ephemeral creek, thus being more susceptible to hydrochemical variations, either due to changes in pond level or by water input from the creek. In fact, the ENSO-controlled variations in rainfall cause significant salinity changes in hypersaline ponds of Kiritimati (see ‘General setting and materials’ section, above), and in particular in Lake 22, which shows the strongest salinity variation (+118‰ in the period from 2002 (132‰, Arp et al., 2012) to 2011 (250‰, Shen et al., 2018), if compared with Lake 2 (+28‰; Saenger et al., 2006; Arp et al., 2012; Shen et al., 2020) and even with the immediately adjacent lake 21 (+57‰; Saenger et al., 2006; Arp et al., 2012; Ionescu et al., 2015).

## Concluding remarks and implications

In summary, the formation mechanism of the Kiritimati microbial mat ooids is interpreted here as product of long-term (~1000 year-scale) mat evolution, especially of certain layers, through a combination of: a) a not too advanced EPS degradation that allows some degree of inhibition of precipitation, hindering or slowing micrite formation (cf. Arp et al., 2012); and b) periodic and significant variations in supersaturation within the mat microenvironment, probably triggered by climate-driven hydrochemical changes in the hypersaline pond, although metabolic changes are also likely to influence local microenvironmental variations during the burial evolution of particular mat layers. Future work on environmental and microbiochemical monitoring of Kiritimati mats or other similar examples, and laboratory experiments replicating these conditions could further refine this interpretation. Nonetheless, this first description of the Kiritimati ooids provides definitive proof that ooids can grow statically within benthic microbial mats, controlled both by biological and environmental factors, a mechanism rarely described (Friedmann et al., 1973; Krumbein, 1983; Gerdes et al., 1994) and poorly clarified. Thus, this study means a significant step forward in the understanding of ooids not merely as physicochemical precipitates, but as particles whose growing mechanism is at least influenced, if not directly controlled, by biotic factors (see a recent review and discussion in Diaz and Eberli, 2019), a hypothesis with increasing evidence that supports it not only from hypersaline environments, but also from freshwater settings (e.g. Wilkinson et al., 1980; Plee et al., 2008; Pacton et al., 2012), normal marine waters (e.g. Diaz et al., 2017; Batchelor et al., 2018; Mariotti et al., 2018) and even from laboratory experiments (e.g. Brehm et al., 2004; 2006). Finally, the Kiritimati ooids demonstrate that care must be taken when interpreting the origin of fossil ooids, one of the oldest (e.g. Siahi et al., 2017; Flannery et al., 2019) and most extensively studied particles of the geological record, providing a modern analogue that will proof useful in explaining fossil ooids with features compatible with an origin associated with benthic microbial communities (e.g. Kalkowsky, 1908; Krumbein, 1983; Neuweiler, 1993; Li et al., 2017; Antoshkina et al., 2020; Zwicker et al., 2020).

## Acknowledgements

This study was funded by a by a postdoctoral “Humboldt Research Fellowship” of the Alexander von Humboldt Foundation. Previous field expeditions and studies were funded by DFG projects FOR 571 (Geobiology of Biofilms) and Re 665/18-2 (Evolution of Biomineralisation). Technical assistant was expertly provided by Dorothea Hause-Reitner with the electron microscope, and by Birgit Röring, Wolfgang Dröse and Axel Hackmann with the laboratory preparation of samples.

## Declarations

### Availability of data and material

All data used for this research are available in the text and figures of the manuscript. The material used is available at the Geobiology Department of the University of Göttingen.

### Code availability

Not applicable

## References

Antoshkina, A.I., E.A. Zhegallo, and S.I. Isaenko. 2020. Microbially mediated organomineralization in Paleozoic carbonate ooids. Paleontological Journal 54: 825–834. https://doi.org/10.1134/S003103012008002X.

Arp, G., J. Hofmann, and J. Reitner. 1998. Microbial Fabric Formation in Spring Mounds (“Microbialites”) of Alkaline Salt Lakes in the Badain Jaran Sand Sea, PR China. PALAIOS 13: 581–592, https://doi.org/10.2307/3515349.

Arp, G., A. Reimer, and J. Reitner. 1999. Calcification in cyanobacterial biofilms of alkaline salt lakes, European Journal of Phycology 34: 393–403, https://doi.org/10.1080/09670269910001736452.

Arp, G., G. Helms, K. Karlinska, G. Schumann, A. Reimer, J. Reitner, and J. Trichet. 2012. Photosynthesis versus Exopolymer Degradation in the Formation of Microbialites on the Atoll of Kiritimati, Republic of Kiribati, Central Pacific. Geomicrobiology Journal 29: 29–65, https://doi.org/10.1080/01490451.2010.521436, 2012.

Bates, R.L, and J.A. Jackson. 1980. Glossary of Geology, 2^nd^ edition. Falls Church, Virginia: American Geological Institute.

Bathurst, R.G.C. 1968. Precipitation of ooids and other aragonite fabrics in warm seas. In Recent Developments in Carbonate Sedimentology in Central Europe, ed. G. Muller and G.M. Friedman, pp. 1–10. Springer, Berlin.

Baumgartner, L.K., R.P. Reid, C. Dupraz, A.W. Decho, D.H. Buckley, J.R. Spear, K.M. Przekop, and P.T. Visscher. 2006. Sulfate reducing bacteria in microbial mats: changing paradigms, new discoveries. Sedimentary Geology 185: 131–145. https://doi.org/10.1016/j.sedgeo.2005.12.008

Blumenberg, M., V. Thiel, and J. Reitner. 2015. Organic matter preservation in the carbonate matrix of a recent microbial mat – Is there a ‘mat seal effect’? Organic Geochemistry 87: 25–34, https://doi.org/10.1016/j.orggeochem.2015.07.005.

Brehm, U., K.A. Palinska, and W.E. Krumbein. 2004. Laboratory cultures of calcifying biomicrospheres generate ooids - A contribution to the origin of oolites. Carnets de Géologie /Notebooks on Geology, Letter 2004/03. http://paleopolis.rediris.es/cg_archives/04L03/

Brehm, U., W.E. Krumbein, and K.A. Palinska. 2006. Biomicrospheres Generate Ooids in the Laboratory. Geomicrobiology Journal 23: 545–550. https://doi.org/10.1080/01490450600897302.

Bucher, W.H. 1918. On oölites and spherulites. The Journal of Geology 26: 583–609. https://doi.org/10.1086/622622

Burne, R.V., J.C. Eade, and J. Paul. 2012. The Natural History of Ooliths: Franz Ernst Brückmann’s Treatise of 1721 and its Significance for the Understanding of Oolites. Hallesches Jahrbuch für Geowissenschaften 34: 93–114.

Chen, M., Conroy, J.L., and Fouke, B.W. 2020. The distribution of organic matter in aragonite spherules: Insights into potential biotic mechanisms of spherule formation. GSA Abstracts with Programs 52 (6), Paper No. 214–7. https://doi.org/10.1130/abs/2020AM-353002.

Chidsey, T.C., M.D. Vanden Berg, and D.E. Eby. 2015. Petrography and characterization of microbial carbonates and associated facies from modern Great Salt Lake and Uinta Basin’s Eocene Green River Formation in Utah, USA. In Microbial Carbonates in Space and Time, ed. D.W.J. Bosence et al., 261–286. Geological Society of London, Special Publication 418.

Davies, P.J., B. Bubela, and J. Ferguson. 1978. The formation of ooids. Sedimentology 25: 703–730. https://doi.org/10.1111/j.1365-3091.1978.tb00326.x

Défarge, C., J. Trichet, A.M. Jaunet, M. Robert, J. Tribble, and F.J. Sansone. 1996. Texture of microbial sediments revealed by cryo-scanning electron microscopy. Journal of Sedimentary Research 66: 935–947. https://doi.org/10.1306/D4268446-2B26-11D7-8648000102C1865D.

Diaz, M.R., and G.P. Eberli. 2019. Decoding the mechanism of formation in marine ooids: A review. Earth-Science Reviews 190: 536–556. https://doi.org/10.1016/j.earscirev.2018.12.016

Diaz, M.R., G.P. Eberli, P. Blackwelder, B. Phillips, and P.K. Swart. 2017. Microbially mediated organomineralization in the formation of ooids. Geology 45: 771–774. https://doi.org/10.1130/G39159.1.

Duguid, S.M.A., T.K. Kyser, N.P. James, and E.C. Rankey. 2010. Microbes and ooids. Journal of Sedimentary Research 80: 236–251. https://doi.org/10.2110/jsr.2010.027.

Eardley, A.J. 1938. Sediments of Great Salt Lake, Utah. AAPG Bulletin 22: 1305–1411.

Fabricius, F.H. 1977. Origin of marine oöids and grapestones. Contributions to Sedimentology 7: 113 pp.

Flannery, D.T., A.C. Allwood, R. Hodyss, R.E. Summons, M. Tuite, M.R. Walter, and K.H. Williford. 2019. Microbially influenced formation of Neoarchean ooids. Geobiology 17: 151–160. https://doi.org/10.1111/gbi.12321.

Friedmann, G.M., A.J. Amiel, M. Braun, and D.S. Miller. 1973. Generation of carbonate particles and laminites in algal mats – Example from sea-marginal hypersaline pool, Gulf of Aqaba, Red Sea. AAPG Bulletin 57: 541–557. https://doi.org/10.1306/819A4302-16C5-11D7-8645000102C1865D

Friedmann, G.M., A. Sneh, and R.W. Owen. 1985. The Ras Muhammad Pool: Implications for the Gavish Sabkha. In Hypersaline Ecosystems - The Gavish Sabkha, eds. W.E. Krumbein and G.M. Friedman, 218–237. Springer-Verlag, Berlin.

Gerdes, G., K. Dunajtschik-Piewak, H. Riege, A.G. Taher, W.E. Krumbein, and H.E. Reineck. Structural diversity of biogenic carbonate particles in microbial mats. Sedimentology 41: 1273–1294. https://doi.org/10.1111/j.1365-3091.1994.tb01453.x

Gerdes, G., W.E. Krumbein, and N. Noffke. 2000. Evaporite microbial sediments. In Microbial Sediments, eds. R.E. Riding and S.M. Awramik, 196–208. Springer-Verlag, Berlin. https://doi.org/10.1007/978-3-662-04036-2_22.

Helfrich, P., J. Ball, A. Berger, P. Bienfang, S.A. Cattell, N. Foster, G. Fredholm, B. Gallagher, E. Guinther, G. Krasnick, M. Rakowicz, and M. Valencia. 1973. The feasibility of brine shrimp production on Christmas Island. Sea Grant Technical Report UNIHI-SEAGRANT-TR-73-02, 173 pp.

Hubert, H.L., E.C. Rankey, and C. Omelon. 2018. Organic matter, textures, and pore attributes of hypersaline lacustrine microbial deposits (Holocene, Bahamas). Journal of Sedimentary Research 88: 827–849. http://dx.doi.org/10.2110/jsr.2018.42

Ionescu, D., S. Spitzer, A. Reimer, D. Schneider, R. Daniel, J. Reitner, D. de Beer, and G. Arp. 2015. Calcium dynamics in microbialite-forming exopolymer-rich mats on the atoll of Kiritimati, Republic of Kiribati, Central Pacific. Geobiology 13: 170–180, https://doi.org/10.1111/gbi.12120.

Kahle, C.F. 1974. Ooids from Great Salt Lake, Utah, as an analogue for the genesis and diagenesis of ooids in marine limestones. Journal of Sedimentary Petrology 44: 30–39. https://doi.org/10.1306/74D7296E-2B21-11D7-8648000102C1865D

Kalkowsky, E. 1908. Oolith und Stromatolith im norddeutschen Buntsandstein. Zeitschrift der Deutschen Geologischen Gesellschaft 60: 68–125.

Krumbein, W.E. 1983. Stromatolites – The challenge of a term in space and time. Precambrian Research 20: 493–531. https://doi.org/10.1016/0301-9268(83)90087-6

Krumbein, W.E., and Y. Cohen. 1974. Biogene, klastische und evaporitische Sedimentation in einem mesothermen monomiktischen ufernahen See (Golf von Aqaba). Geologische Rundschau 63: 1035–1065. https://doi.org/10.1007/BF01821322

Li, F., J. Yan, R.V. Burne, Z.Q. Chen, T.J. Algeo, W. Zhang, L. Tian, Y. Gan, K. Liu, and S. Xie. 2017. Paleo-seawater REE compositions and microbial signatures preserved in laminae of Lower Triassic ooids. Palaeogeography, Palaeoclimatology, Palaeoecology 486: 96–107. https://doi.org/10.1016/j.palaeo.2017.04.005

Mariotti, G., S.B. Pruss, R.E. Summons, S.A. Newman, and T. Bosak. 2018. Contribution of Benthic Processes to the Growth of Ooids on a Low-Energy Shore in Cat Island, The Bahamas. Minerals 8: 252. https://doi.org/10.3390/min8060252.

Mikutta, R., M. Kleber, K. Kaiser, and R. Jahn. 2005. Review: Organic matter removal from soils using hydrogen peroxide, sodium hypochlorite, and disodium peroxodisulfate, Soil Science Society of America 69: 120–135, https://doi.org/10.2136/sssaj2005.0120.

Mitterer, R.M. 1968. Amino acid composition of organic matrix in calcareous oolites. Science 162: 1498–1499. https://doi.org/10.1126/science.162.3861.1498

Morrison, R.J., and C.D. Woodroffe. 2009. The soils of Kiritimati (Christmas) Island, Kiribati, Central Pacific: New information and comparison with previous studies. Pacific Science 60: 397–411. https://doi.org/10.2984/049.063.0308

Neuweiler, F. 1993. Development of Albian microbialites and microbialite reefs at marginal platform areas of the Vasco-Cantabrian Basin (Soba Reef area, Cantabria, N. Spain). Facies 29: 213–250. https://doi.org/10.1007/BF02536930

Pacton, M., D. Ariztegui, D. Wacey, M.R. Kilburn, C. Rollion-Bard, R. Farah, and C. Vasconcelos. 2012. Going nano: A new step toward understanding the processes governing freshwater ooid formation. Geology 40: 547–550. https://doi.org/10.1130/G32846.1.

Peryt, T.M. 1983. Classification of coated grains. In Coated Grains, ed. T.M. Peryt, 3–6. Berlin: Springer-Verlag.

Plee, K., D. Ariztegui, R. Martin, and E. Davaud. 2008. Unravelling the microbial role in ooid formation – results of an in situ experiment in modern freshwater Lake Geneva in Switzerland. Geobiology 6: 341–350. https://doi.org/10.1111/j.1472-4669.2007.00140.x.

Reitner, J., P. Gautret, F. Marin, F. Neuweiler. 1995. Automicrites in a modern marine microbialite. Formation model via organic matrices (Lizard Island, Great Barrier Reef, Australia). Bulletin de I’Institut Océanographique, Monaco 14: 237–263.

Reitner, J., G. Arp, V. Thiel, P. Gautret, U. Galling, and W. Michaelis. 1997. Organic matter in Great Salt Lake ooids (Utah, USA) – First approach to a formation via organic matrices. Facies 36: 210–219.

Reitner, J., J. Peckmann, M. Blumenberg, W. Michaelis, A. Reimer, and V. Thiel. 2005. Concretionary methane-seep carbonates and associated microbial communities in Black Sea sediments. Palaeogeography, Palaeoclimatology, Palaeoecology 227: 18–30. https://doi.org/10.1016/j.palaeo.2005.04.033

Richter, D.K. 1983a. Calcareous ooids: A synopsis. In Coated Grains, ed. T.M. Peryt, 71–99. Berlin: Springer-Verlag.

Richter, D.K. 1983b. Classification of coated grains: Discussion. In Coated Grains, ed. T.M. Peryt, 7–8. Berlin: Springer-Verlag.

Sachs, J.P., D. Sachse, R.H. Smittenberg, Z. Zhang, D.S. Battisti, and S. Golubic. 2009. Southward movement of the Pacific intertropical convergence zone AD 1400-1850. Nature Geoscience 2: 519–525. https://doi.org/10.1038/ngeo554

Saenger, C., M. Miller, R.H. Smittenberg, and J.P. Sachs. 2006. A physico-chemical survey of inland lakes and saline ponds: Christmas Island (Kiritimati) and Washington (Teraina) Islands, Republic of Kiribati. Saline Systems 2: 1–15. https://doi.org/10.1186/1746-1448-2-8, 2006.

Schmitt, S., J.L. Conroy, T.M. Flynn, R.A. Sanford, M.C. Higley, M. Chen, and B.W. Fouke. 2019. Salinity, microbe and carbonate mineral relationships in brackish and hypersaline lake sediments: A case study from the tropical Pacific coral atoll of Kiritimati. The Depositional Record 5: 212–229. https://doi.org/10.1002/dep2.71.

Schneider, D., G. Arp, A. Reimer, J. Reitner, and R. Daniel. 2013. Phylogenetic analysis of a microbialite-forming microbial mat from a hypersaline lake of the Kiritimati Atoll, Central Pacific. PLoS ONE 8: e66662. https://doi.org/10.1371/journal.pone.0066662

Schoonmaker, J., G.W. Tribble, S.V. Smith, and F.T. Mackenzie. 1985. Geochemistry of saline ponds, Kiritimati (Republic of Kiribati). Proceedings of the 5th International Coral Reef Congress, Vol. 3, pp. 439–444.

Shen, Y., V. Thiel, J.P. Duda, and J. Reitner. 2018. Tracing the fate of steroids through a hypersaline microbial mat (Kiritimati, Kiribati/Central Pacific). Geobiology 16: 307–318, https://doi.org/10.1111/gbi.12279.

Shen, Y., V. Thiel, P. Suarez-Gonzalez, S.W. Rampen, and J. Reitner. 2020. Sterol preservation in hypersaline microbial mats. Biogeosciences 17: 649–666, https://doi.org/10.5194/bg-17-649-2020.

Shtukenberg, A.G., Y.O. Punin, E. Gunn, and B. Kahr. 2012. Spherulites. Chemical Reviews 112: 1805–1838. https://doi.org/10.1021/cr200297f.

Siahi, M., A. Hofmann, S. Master, C.W. Mueller, and A. Gerdes. 2017. Carbonate ooids of the Mesoarchaean Pongola Supergroup, South Africa. Geobiology 15: 750–766. https://doi.org/10.1111/gbi.12249.

Simone, L. 1981. Ooids: A review. Earth-Science Reviews 16: 319–355. https://doi.org/10.1016/0012-8252(80)90053-7

Suarez-Gonzalez, P., D. Hause-Reitner, Y. Shen, N. Schäfer, and J. Reitner. 2017. The makings of a microbialite – Insights into the earliest stages of mineralization within microbial mats (Kiritimati Island, Central Pacific). 4^th^ International Conference of Geobiology, Wuhan, China. Abstract book, pp. 134–135.

Suarez-Gonzalez, P., M.I. Benito, I.E. Quijada, R. Mas, S. Campos-Soto. 2019. ‘Trapping and binding’: A review of the factors controlling the development of fossil agglutinated microbialites and their distribution in space and time. Earth-Science Reviews 194: 182–215. https://doi.org/10.1016/j.earscirev.2019.05.007

Suess, E., and D. Fütterer. 1972. Aragonitic ooids: experimental precipitation from seawater in the presence of humic acids. Sedimentology 19: 129–139. https://doi.org/10.1111/j.1365-3091.1972.tb00240.x

Teichert, C. 1970. Oolite, oolith, ooid: Discussion. AAPG Bulletin 54: 1748–1749.

Trichet, J., and C. Défarge. 1995. Non-biologically supported organomineralization. Bulletin de I’Institut Océanographique, Monaco 14: 203–236.

Trichet, J., C. Defarge, J. Tribble, G. Tribble, and F. Sansone. 2001. Christmas Island lagoonal lakes, models for the deposition of carbonate-evaporite-organic laminated sediments. Sedimentary Geology 140: 177189. https://doi.org/10.1016/S0037-0738(00)00177-9, 2001.

Trower, E.J., M.D. Cantine, M.L. Gomes, J.P. Grotzinger, A.H. Knoll, M.P. Lamb, U. Lingappa, S.S. O’Reilly, T.M. Present, N. Stein, J.V. Strauss, and W.W. Fischer. 2018. Active ooid growth driven by sediment transport in a high-energy shoal, Little Ambergris Cay, Turks and Caicos Islands. Journal of Sedimentary Research 88: 1132–1151. http://dx.doi.org/10.2110/jsr.2018.59.

Valencia, M.J (1977) Christmas Island (Pacific Ocean): Reconnaissance geologic observations. Atoll Research Bulletin 197: 1–14.

Verrechia, E.P., P. Freytet, K.E. Verrechia, and J.L. Dumont. 1995. Spherulites in calcrete laminar crusts: Biogenic CaCO3 precipitation as a major contributor to crust formation. Journal of Sedimentary Research A65: 690–700. https://doi.org/10.1306/D426819E-2B26-11D7-8648000102C1865D

Weber, G.W. 2014. Another link between archaeology and anthropology: Virtual anthropology. Digital Applications in Archaeology and Cultural Heritage 1: 3–11. http://dx.doi.org/10.1016/j.daach.2013.04.001.

Wilkinson, B.H., B.N. Pope, and R.M. Owen. 1980. Nearshore ooid formation in a modern temperate region marl lake. Journal of Geology 88: 697–704.

Wilkinson, B.H., R.M. Owen, and A.R. Carroll. 1985. Submarine hydrothermal weathering, global eustasy, and carbonate polymorphism in Phanerozoic marine oolites. Journal of Sedimentary Petrology 55: 171–183.

Zwicker, J., D. Smrzka, F. Steindl, M.E. Böttcher, E. Libowitzky, S. Kiel, and J. Peckmann. 2020. Mineral authigenesis within chemosynthetic microbial mats: Coated grain formation and phosphogenesis at a Cretaceous hydrocarbon seep, New Zealand. The Depositional Record. https://doi.org/10.1002/dep2.123

